# The use of two-sample methods for Mendelian randomization analyses on single large datasets

**DOI:** 10.1101/2020.05.07.082206

**Authors:** Cosetta Minelli, M. Fabiola Del Greco, Diana A. van der Plaat, Jack Bowden, Nuala A. Sheehan, John Thompson

## Abstract

**Background:** With genome-wide association data for many exposures and outcomes now available from large biobanks, one-sample Mendelian randomization (MR) is increasingly used to investigate causal relationships. Many robust MR methods are available to address pleiotropy, but these assume independence between the gene-exposure and gene-outcome association estimates. Unlike in two-sample MR, in one-sample MR the two estimates are obtained from the same individuals, and the assumption of independence does not hold in the presence of confounding.

**Methods:** With simulations mimicking a typical study in UK Biobank we assessed the performance, in terms of bias and precision of the MR estimate, of the fixed-effect and (multiplicative) random-effects meta-analysis method, weighted median estimator, weighted mode estimator and MR-Egger regression, used in both one-sample and two-sample data. We considered scenarios differing for: presence/absence of a true causal effect; amount of confounding; presence and type of pleiotropy (none, balanced or directional).

**Results:** Even in the presence of substantial correlation due to confounding, all methods performed well when used in one-sample MR except for MR-Egger, which resulted in bias reflecting direction and magnitude of the confounding. Such bias was much reduced in the presence of very high variability in instrumental strength across variants (I^2^_GX_ of 97%).

**Conclusions:** Two-sample MR methods can be safely used for one-sample MR performed within large biobanks, expect for MR-Egger. MR-Egger is not recommended for one-sample MR unless the correlation between the gene-exposure and gene-outcome estimates due to confounding can be kept low, or the variability in instrumental strength is very high.

**Key Messages:**

- Current availability of phenotypic and genetic data from large biobanks, such as UK Biobank, has led to increasing use of one-sample Mendelian randomization (MR) to investigate causal relationships in epidemiological research
- Robust MR methods have been developed to address pleiotropy, but they assume independence between the gene-exposure and gene-outcome association estimates; this holds in two-sample MR but not in one-sample MR
- We illustrate the practical implications, in terms of bias and precision of the MR causal effect estimate, of using robust two-sample methods in one-sample MR studies performed within large biobanks
- Two-sample MR methods can be safely used for one-sample MR performed within large biobanks, expect for MR-Egger regression
- MR-Egger is not recommended for one-sample MR unless the correlation between the gene-exposure and gene-outcome estimates due to confounding can be kept low, or the variability in instrumental strength is very high

## Introduction

Mendelian randomization (MR) is an instrumental variable approach to investigate the causal effect of an exposure on an outcome by using genetic variants, typically single nucleotide polymorphisms (SNPs), as instruments for the exposure^1^; the causal effect is indirectly estimated from the gene-exposure and gene-outcome associations. The validity of MR relies on instrumental variable assumptions^2^, the most problematic being the absence of pleiotropy whereby the genetic instruments modify the outcome only through the exposure of interest and not through any other independent pathway.

When full genetic, exposure and continuous outcome data are all available within the same study (“one-sample MR”), the causal effect can be estimated using the two-stage least-square (2SLS) method; for a binary outcome, the analogue is a two-stage estimator with a logistic or log-linear model at the second-stage (exposure–outcome) regression^3^. When only summary statistics (β coefficients and standard errors) for gene-exposure and gene-outcome associations are available from separate studies (“two-sample MR”), the causal effect is often estimated by first deriving SNP-specific causal effect estimates as the gene-outcome estimate divided by the gene-exposure estimate (Wald estimator), and then pooling them using inverse-variance weighted fixed-effect meta-analysis (IVW FE)^4, 5^. Two-sample MR has been widely used to exploit summary data from large genetic consortia^5^, and this has addressed the issue of low statistical power typical of MR^6^. However, the two-sample approach has the important limitation of having to assume that the samples are homogeneous, so that the gene-exposure associations are identical across the samples. This may be violated in practice^7^. Recently, large population-based biobanks have made available individual-level genome-wide data and data on a variety of exposures and outcomes, thus allowing well-powered one-sample MR studies. An important example is the UK Biobank (UKB), where individual-level data are publicly available for about 500,000 individuals aged 40-69^8^.

While both the 2SLS method for one-sample MR and the IVW FE method for two-sample MR assume no pleiotropy, several alternative methods that are robust to pleiotropy have been developed for two-sample MR^9^. It is therefore tempting to use robust two-sample MR methods in a one-sample MR. The problem is that these methods assume independence between the gene-exposure and gene-outcome estimates, as would be the case when they are obtained from separate non-overlapping samples^10^. This assumption does not hold in one-sample MR, where the two estimates are obtained from the same individuals and are therefore correlated.

Using simulations, we investigated the practical implications of applying methods that assume independence between the gene-exposure and gene-outcome estimates (referred to as “two-sample MR methods”) to a one-sample MR in the specific context of a large biobank, focusing on a continuous outcome. We considered the case where the SNP discovery for the one-sample MR is based on evidence from previous studies, thus avoiding any issue with the winner’s curse^11^. To inform our simulations we reviewed 10 published studies that used two-sample MR methods in one-sample MR within UKB. In the simulations we assessed the performance, in terms of bias and precision of the causal effect estimate, of five different two-sample MR methods (fixed-effect and multiplicative random-effects meta-analysis, weighted median estimator, weighted mode estimator, and MR-Egger regression), in both one-sample and two-sample data. We considered scenarios in which there was a true causal effect or not, in which the amount of confounding varied (implicit in the varying amount of correlation between exposure and outcome errors), and in which pleiotropy varied (none, balanced or directional).

## Methods

### Examples of published UKB studies

To inform our simulations, we searched PubMed on 25 April 2019 to identify examples of one-sample MR studies performed within UKB (“*UK Biobank[Title/Abstract] AND (Mendelian randomization[Title/Abstract] OR Mendelian randomisation[Title/Abstract])*”). We reviewed the 10 most recent studies identified that had used two-sample methods with multiple instruments. From these we extracted information on sample size, number of instruments, total variance of the exposure explained, F statistic, and MR methods used in main and in secondary analyses. For studies using MR-Egger, we also recorded the variability in instrument strength, expressed as heterogeneity in gene-exposure estimates across SNPs (I^2^_GX_), since we have previously shown that MR-Egger works well only when this is large, with recommended I^2^_GX_ over 90%^12^. Although 110 eligible MR investigations were reported in the 10 papers, for each paper we only considered those on different exposures (different sets of instruments) or population subgroups (different sample sizes). This resulted in 27 MR investigations, whose characteristics are summarised in Supplementary Table 1. The sample size varied from 180,957 to 376,435 (median: 318,664), the number of instruments from 2 to 520 (median: 68), and the variance explained, reported in 5 studies, from 0.2% to 7.3% (median: 1.8%). Only 3 studies reported the F statistic; of these, only one study (reporting on 4 MR investigations) provided values for all SNPs, which varied from 10.1 to 382.5.

The most commonly used two-sample methods were: MR-Egger regression (N=10 studies); weighted median estimator (N=9); IVW (N=8), mostly as the main analysis, with only 3 studies specifying whether a fixed-effect or a random-effects model was used; weighted mode estimator (N=3). All studies used MR-Egger but none reported an I^2^_GX_ value, although one mentioned the limited variability in instrument strengths as an explanation for the limited power of the MR-Egger analysis^13^. A one-sample MR method was also used in 3 studies: 2SLS (N=2) and a maximum likelihood method3 (N=1).

### Methods for the simulations

Mirroring a typical analysis using UKB, the simulations created data on 300,000 individuals and 100 independent SNPs with allele frequencies between 1% and 99%. Both the exposure, X, and the outcome, Y, were continuous with normally distributed errors, and all relationships between X, Y and the genotype, G, were linear with no interactions. SNP coefficients for the G-X association were simulated using an exponential distribution, resulting in many SNPs with small effects and a few with large effects (Supplementary Methods). The individual SNPs had an average F-statistic of 67.9; only 5% of the SNPs had an F statistic below 10 and would generally be considered as weak ^14^. On average, the 100 SNPs explained 2.3% of the variance in X. The average I^2^_GX_ was 91%.

We simulated data with no causal effect of X on the outcome, Y, and data where the causal effect was 1.0. A causal effect of 1.0 was strong enough that the 100 SNPs explained 0.2% of the variance of the outcome in the absence of pleiotropy. For each scenario, different degrees of confounding between X and Y were simulated by generating a correlation between the error components in X and Y of −0.4, −0.2, 0, 0.2, 0.4 (a correlation of 0 represents no confounding).

As well as the situation of no pleiotropy, we simulated data in which 20% of the SNPs were pleiotropic (details in Supplementary Methods). The pleiotropic effects were generated from the same distribution as the G-X effects but independently, so that the InSIDE (Instrument Strength Independent of Direct Effect) assumption needed for two-sample methods holds^15^. For balanced pleiotropy, the pleiotropic effects were given a random sign so that the average was zero, while for directional pleiotropy, the effects were all positive.

For comparison, genuine two-sample data were also simulated by creating two datasets under identical conditions and taking the G-X estimates from one and the G-Y estimates from the other.

All simulations were run for 1,000 times. Further details of the simulation parameters are reported in Supplementary Methods.

### MR methods compared

The two-sample methods investigated were IVW FE^5^ and four methods robust to pleiotropy, which are based on different assumptions about its nature: multiplicative random-effects meta-analysis (IVW RE), that has been recommended over the additive RE model (see Supplementary Methods)^16^; weighted median estimator^17^; weighted mode estimator^18^; MR-Egger regression^15^. In the one-sample MR we also calculated the 2SLS estimator^3^, which represents the gold standard in the absence of pleiotropy. Further details are reported in Supplementary Methods.

All two-sample methods were implemented using the MendelianRandomization R package (https://CRAN.R-project.org/package=MendelianRandomization), while for 2SLS we used the AER R package (https://CRAN.R-project.org/package=AER).

## Results

Overall, the results of our simulations show that, except for MR-Egger, two-sample methods perform similarly when applied to a one-sample MR with a large sample size (300,000 in our simulations) or a two-sample MR of the same size. In the presence of confounding between X and Y, MR-Egger used in one-sample MR gives biased results that reflect the magnitude and direction of the confounding; in particular, the bias is in the direction of the observational association, which can be viewed as the sum of the causal effect of X on Y, and the confounder effect (see causal diagram in Supplementary Methods).

Figures 1 and 2 show boxplots of the 1,000 MR estimates generated when there is no true causal effect and with a causal effect of 1, respectively. The point estimates for the causal effect from IVW FE, IVW (multiplicative) RE and 2SLS are theoretically identical^19^, as reflected in Figures 1 and 2. Numerical results corresponding to Figures 1 and 2, expressed as mean, standard error, coverage and root mean square error (RMSE) of the causal effect, are reported in Supplementary Tables 2, 3 and 4 for each pleiotropy scenario. To improve readability of the plots, outliers (values more than 0.8 above/below the true value) were removed from the graphs, but not from the results in Supplementary Tables 2-4. Supplementary Table 5 includes the ordinary least squares (OLS) estimate for regression of Y on X in one-sample data, which indicates the amount of confounding present in the simulations.

**Figure 1.**
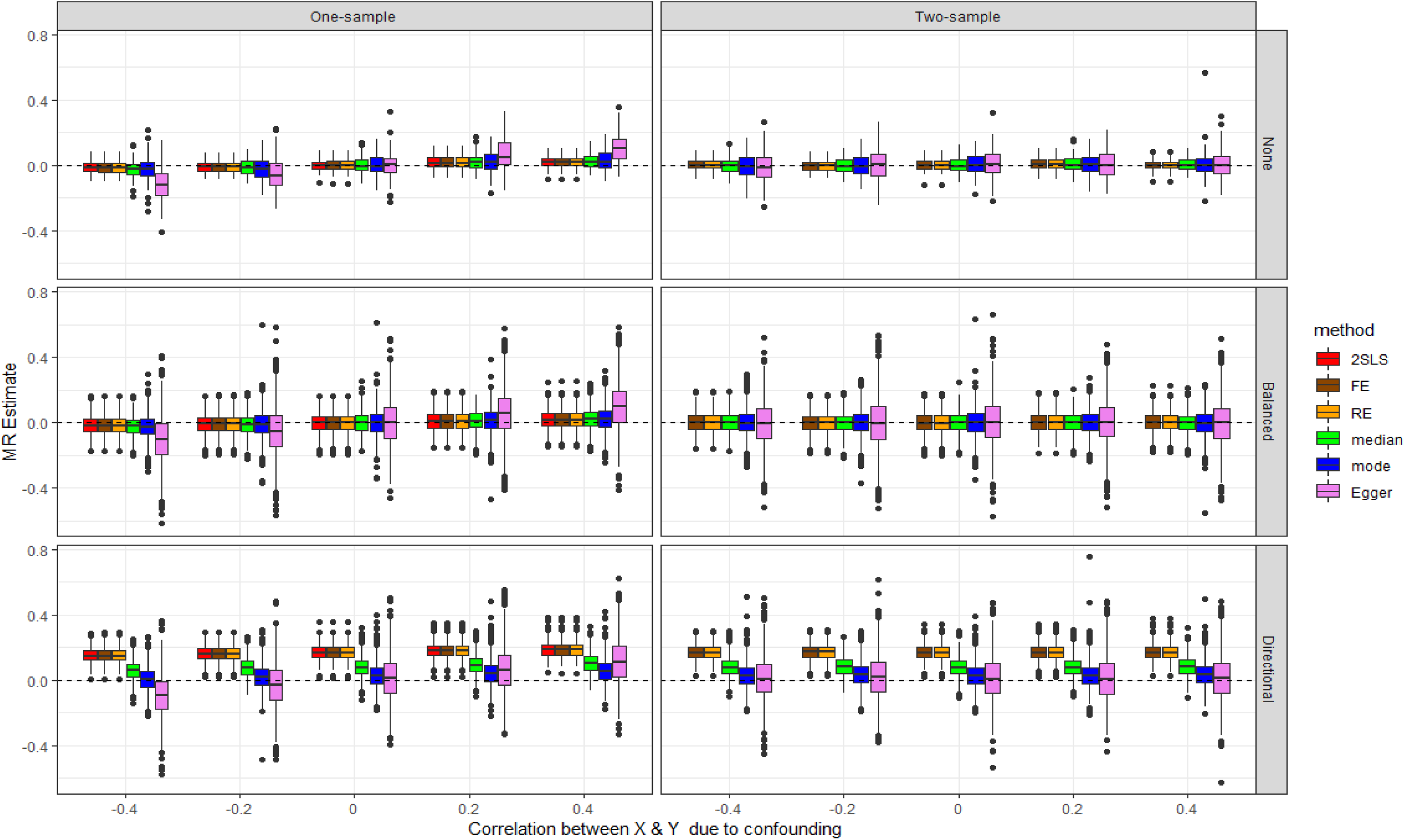
Simulations with no causal effect: Box plots summarising the results of all methods across the 1,000 simulations, in scenarios with no pleiotropy, balanced pleiotropy and directional pleiotropy, and for both one-sample and two-sample MR. Outliers have been removed (see text).

**Figure 2.**
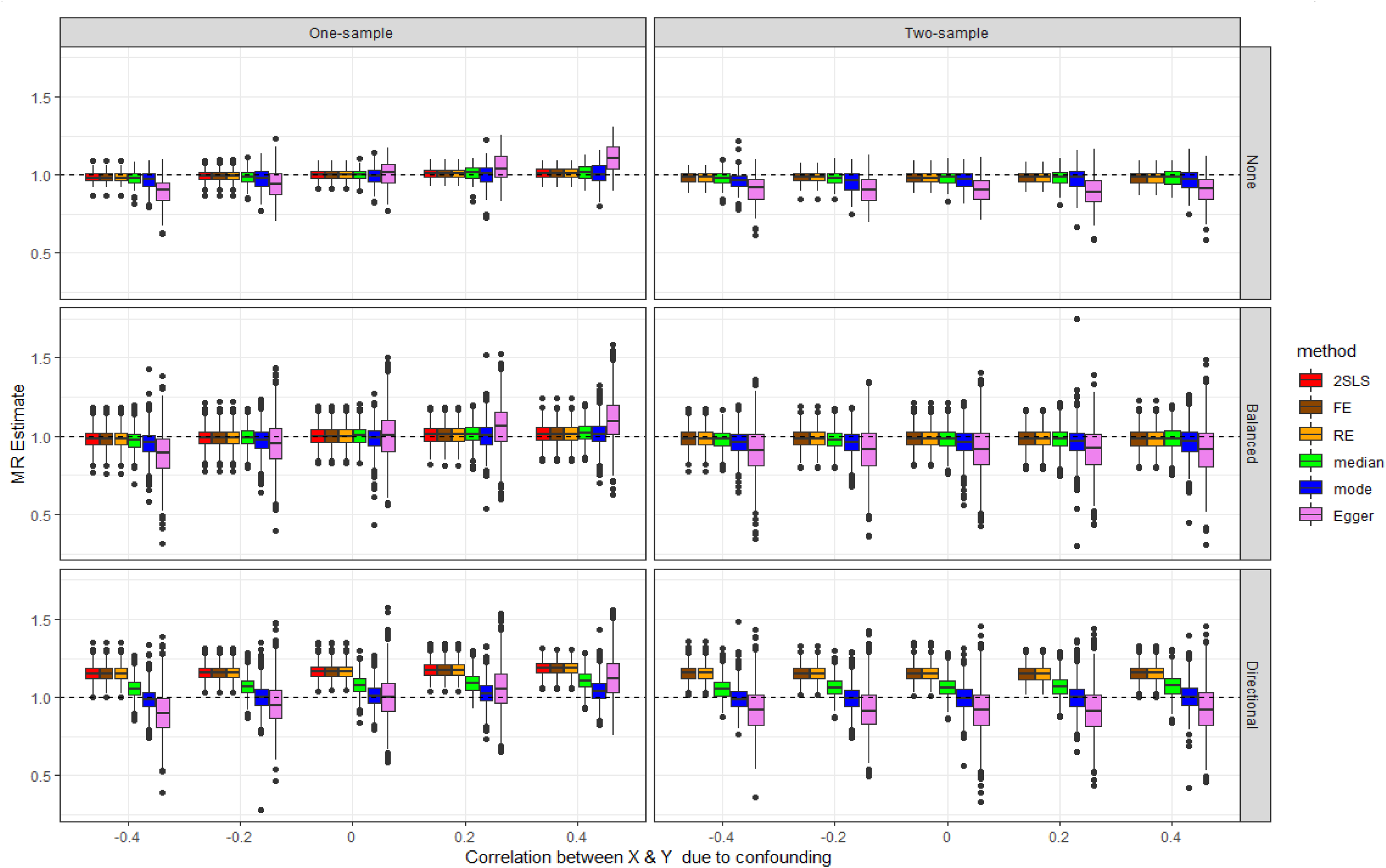
Simulations with a causal effect of 1. Box plots summarising the results of all methods across the 1,000 simulations, in scenarios with no pleiotropy, balanced pleiotropy and directional pleiotropy, and for both one-sample and two-sample MR. Outliers have been removed (see text).

### Scenarios with no pleiotropy

The top panels of Figures 1 and 2 present the results for the scenarios with no pleiotropy.

In the absence of a causal effect (Figure 1), all two-sample methods applied to one-sample MR (left) give the same results on average as when applied to a genuine two-sample MR (right) in the absence of confounding, since the G-X and G-Y estimates become independent even in a one-sample MR. In the presence of confounding, all two-sample methods other than MR-Egger show only minimal bias in the direction of the confounding when applied to one-sample MR; the fact that 2SLS shows the same minimal bias suggests some weak instrument effects affecting all methods. For MR-Egger, however, the bias in the direction of the confounding is substantial. The p-values of MR-Egger analysis are shown in the Q-Q plots of Figure 3 (corresponding Q-Q plots for scenarios with balanced and directional pleiotropy in Supplementary Figure 1). Figure 3 shows increasing departure of the observed from the expected p-values under the null with increasing confounding when MR-Egger is applied to the one-sample MR (left), but not in a two-sample MR (right). Since the points are above expectation, in the presence of confounding, MR-Egger in one-sample MR will mislead by indicating a significant causal effect more often than it should.

**Figure 3.**
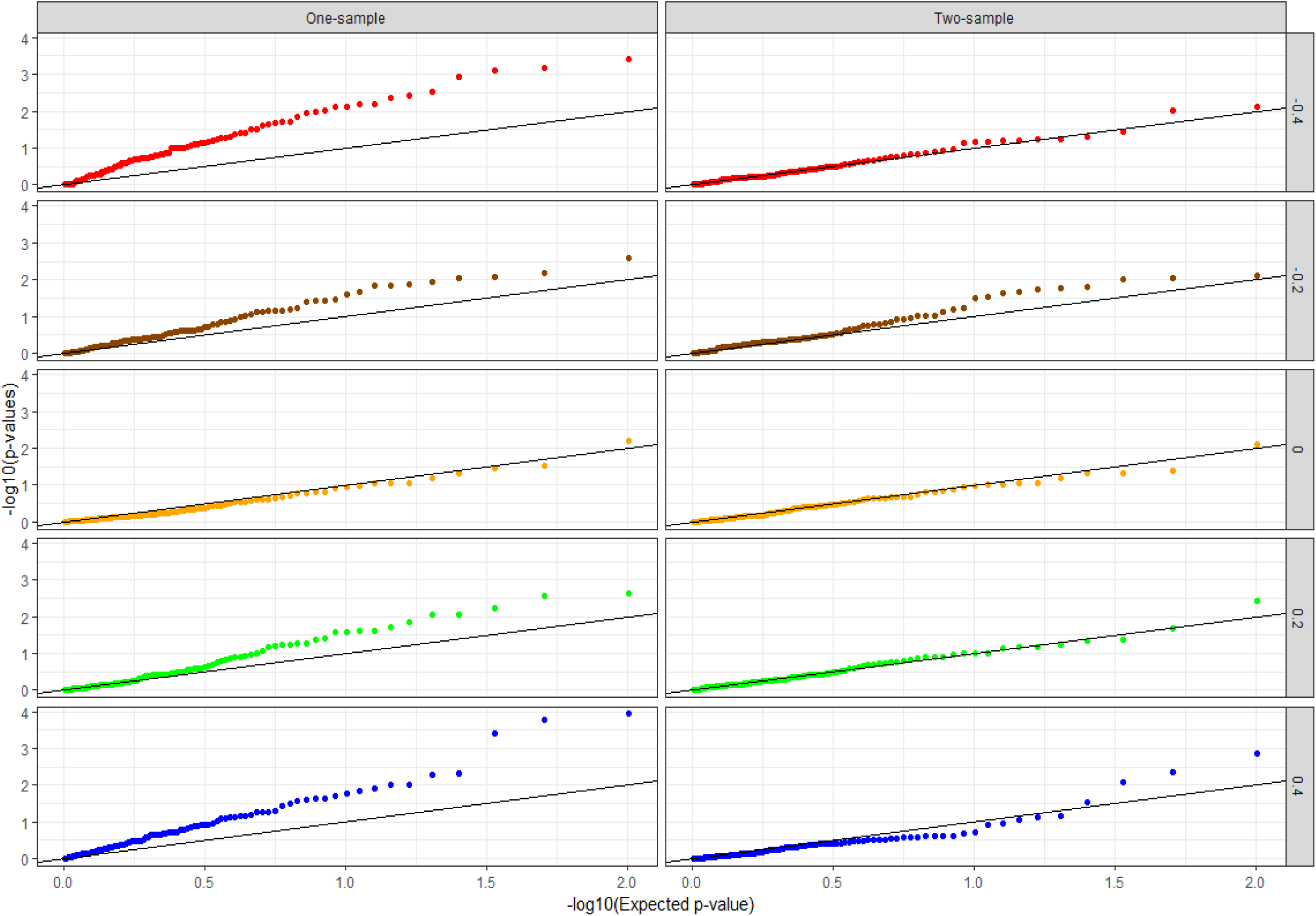
Simulations with no causal effect and no pleiotropy. Q-Q plot of MR-Egger null p-values [plot of -log10(p-value) against its expectation across simulations] for different levels of correlation between X and Y due to confounding, in one-sample and two-sample MR.

Findings for the scenarios with a true causal effect (Figure 2) tend to follow the same pattern as for those with no effect. In the presence of confounding, all two-sample methods used in one-sample MR show a very small bias in the direction of the confounding, except for MR-Egger where this bias is substantial. Interestingly, all methods applied to a genuine two-sample MR show a bias towards the null, but this bias is of very small magnitude expect for MR-Egger where it can be very pronounced. Further investigation (data not shown) indicated that this is due to regression dilution bias, which for MR-Egger is accurately quantified by the I^2^_GX_ statistic that measures the sample variation in instrument strength. This is in line with our previous findings of this bias being much more marked for MR-Egger than for the IVW approach^12^.

To further investigate the behaviour of MR-Egger in the one-sample MR, in a secondary analysis we repeated the simulations after increasing the variability in instrument strength, from a value of I^2^_GX_ of 91% to 97%. The bias in the one-sample MR remained in both scenarios of absence and presence of a causal effect (Supplementary Figure 2a and 2b, respectively), but its magnitude was substantially reduced.

### Scenarios with pleiotropy

When there is balanced pleiotropy, as expected all methods produce more variable causal effect estimates in both one-sample and genuine two-sample MR, and once again MR-Egger gives the most variable estimates. In general, both in the absence and presence of a causal effect (Figures 1 and 2), we observed the same trends as with no pleiotropy, except for the weighted median and weighted mode estimators that were here more similar to the 2SLS, IVW FE and IVW RE methods in terms of variability. 2SLS and IVW FE, which do not allow for pleiotropy, estimated standard errors that were too small resulting in lower coverage and exaggerated significance, while the other methods gave reasonable estimated standard errors and coverage (Supplementary Table 3).

Under directional pleiotropy, as expected the 2SLS, IVW FE and IVW RE methods perform badly in terms of bias across all scenarios (absence/presence of a causal effect; one-sample/two-sample MR; all levels of confounding; Figures 1 and 2). The weighted median estimator tends to be more biased than the weighted mode estimator and MR-Egger across all scenarios. As a consequence of bias accompanied by a small standard error, the coverage for 2SLS, IVW FE and IVW RE is very low (1% to 17% instead of 95% across all one-sample MR scenarios, Supplementary Table 4); for the weighted median estimator, the coverage tends to be worse than that of the weighted mode estimator and MR-Egger. Although in general MR-Egger performs well under directional pleiotropy, the same pattern of bias in the direction of the confounding is observed when applied to the one-sample MR.

## Discussion

With genome-wide association data for many exposures and outcomes becoming readily available from large biobanks such as UK Biobank, one-sample MR is being increasingly used to identify and estimate causal relationships. The main obstacle to the validity of MR is the presence of pleiotropy; several methods robust to pleiotropy are available, but these have been developed for two-sample MR. Our study evaluated the practical implications of using two-sample methods for one-sample MR, where the assumption of independence between the gene-exposure and gene-outcome estimates measured in the same individuals does not hold in the presence of confounding. We show that most two-sample methods, in particular the fixed-effect and (multiplicative) random-effects meta-analysis, the weighted median estimator and the weighted mode estimator, perform well when used in one-sample MR applied to a large sample size, even in the presence of substantial confounding. However, MR-Egger regression does not, with bias in the direction of the confounding increasing with the magnitude of the induced correlation. Our findings confirm previous suggestions of a bias of the MR-Egger estimate towards that of the confounded observational association in the one-sample setting, that has been attributed to the presence of weak instruments to which MR-Egger is particularly vulnerable^20, 21^. MR-Egger may suggest a causal effect when in fact there is none when used in one-sample MR in the presence of confounding, with a positive or negative spurious effect according to the direction of the confounding. Therefore, as opposed to what happens in a two-sample MR, MR-Egger is not necessarily conservative when used in the one-sample MR, since type 1 error is also affected in the presence of confounding. Under directional pleiotropy, although as expected MR-Egger performed better, we observed the same pattern of bias; the weighted mode estimator appeared to be a preferable option in one-sample MR in the scenarios considered in our simulations.

The bias for MR-Egger used in one-sample MR was much attenuated when the method is used at its best, that is with a maximum variability in instrumental strength across variants that can be measured with the I^2^_GX_. However, we show that I^2^_GX_ needs to be much higher than the recommended 90%^12^ in order to reduce the bias in MR-Egger; in our simulations, substantial reduction was obtained by increasing it from 91% to 97%. Increasing the I^2^_GX_ threshold recommended for MR-Egger when used in one-sample MR could be an option, although in practise this may limit the application of the method. An easier option would be to reduce the confounding between exposure and outcome as much as possible, since this would reduce the correlation between the gene-exposure and gene-outcome estimates. In a given one-sample MR study the observed correlation between the two estimates will be induced by both the true causal effect and the confounding; we therefore provide a simple way to disentangle the two and estimate the latter, which is what should be monitored (see derivation of “residual correlation” under Parameters monitored, Supplementary Methods). In practice, the problem with this is that adjustment for potential confounders is not necessarily desirable; in fact, extensive adjustment is not recommended in MR as this may bias estimates if the variable adjusted for represents a mediator (on the causal pathway from the genetic variants to the outcome) or if the adjustment induces collider bias^22^.

The reason why MR-Egger performs worse than the others in a one-sample setting is unclear and requires further investigation. MR-Egger regresses the individual gene-outcome estimates on the gene-exposure estimates. If the estimates are independent, under the InSIDE assumption the slope of the regression line is an unbiased estimate of the causal effect even when pleiotropy is present. In a one-sample MR, the correlation between the gene-outcome and the gene-exposure estimates in the presence of confounding does not only affect the standard error of the MR-Egger estimate, as expected, but also biases the causal estimate in the direction of the confounding.

This study has limitations that need to be addressed with further methodological work. Our findings only apply to the use of two-sample methods in one-sample MR studies performed in large datasets, such as UK Biobank. The problems that we observed are likely to be exacerbated in small samples.

When genetic variants are considered one at a time, as in all two-sample methods, other variants with a pleiotropic effect will induce further confounding between exposure and outcome. While reflected in our simulations, this confounding cannot be disentangled from the other two sources of correlation (classical confounding and causal effect) in any given one-sample MR study; however, such confounding is very likely to be small. Moreover, our simulations for the pleiotropy scenarios assumed independence of the pleiotropic effects from the effects of the instruments on the exposure (InSIDE assumption); in practice, violations of this assumption would also contribute to create a correlation between the gene-exposure and gene-outcome estimates.

In conclusion, fixed-effect and (multiplicative) random-effects meta-analysis, weighted median estimator and weighted mode estimator are two-sample MR methods that can be safely used for one-sample MR studies performed within large biobanks, such as UK Biobank. On the contrary, the use of MR-Egger regression for one-sample MR is not recommended unless the correlation between the gene-exposure and gene-outcome estimates induced by confounding can be kept low, or the variability in instrumental strength across variants is very high. Further work is required to correct for the bias in the MR estimate when MR-Egger is used in the one-sample setting.

## Supplementary Material

**Supplementary Methods.** Details on the parameters used to generate the simulated data, the analyses applied, and the performance measures used.

**Supplementary Figure 1.** Results of the simulations with no causal effect in the presence of pleiotropy: Q-Q plot of MR-Egger null p-values for different levels of correlation between X and Y due to confounding, in one-sample and two-sample MR. a) Balanced pleiotropy; b) Directional pleiotropy.

**Supplementary Figure 2.** Results of secondary analyses where the simulated G-X effect sizes have larger variability (I^2^_GX_ of 0.97). Outliers have been excluded to improve readability of the plots. a) No causal effect; b) Causal effect of 1.

**Supplementary Table 1.** Characteristics and methods used in 27 MR investigations reported in the 10 papers reviewed. We only included one-sample MR analyses performed within UKB; when MR investigations were performed for multiple outcomes, we only considered one outcome per paper.

**Supplementary Tables 2 to 4.** Results of the simulations for: no pleiotropy (Supp. Tab. 2), balanced pleiotropy (Supp. Tab. 3); directional pleiotropy (Supp. Tab. 4). Reported are the causal effect estimate and its standard error, as well as coverage and root mean square error.

**Supplementary Table 5.** Ordinary least squares (OLS) results for the regression analysis of Y on X in the one-sample data. Reported are the causal effect estimate and its standard error; bias; coverage and root mean square error.

