## Supplement for "The use of two-sample methods for Mendelian randomization analyses on single large datasets"

### Supplementary Methods

Here we provide details on our simulation work, in particular on the parameters used to generate the simulated data, the analyses applied, and the performance measures used.

The simulations compare two-sample methods using single sample data and two-sample methods using two-sample data. To do this, we simulate a single sample containing X, Y and G and then using exactly the same parameters we simulate a second sample containing X and G. The first dataset then becomes the Y and G data for the genuine two-sample analysis.

Below is the causal diagram describing the situation our simulations refer to, with G = genetic variant; X = exposure; Y = outcome; U = unmeasured confounders; α = effect of G on X; β = causal effect of X on Y; γ = pleiotropic effect of G on Y. The confounding effect of U was simulated by generating a correlation in the errors of X and Y.

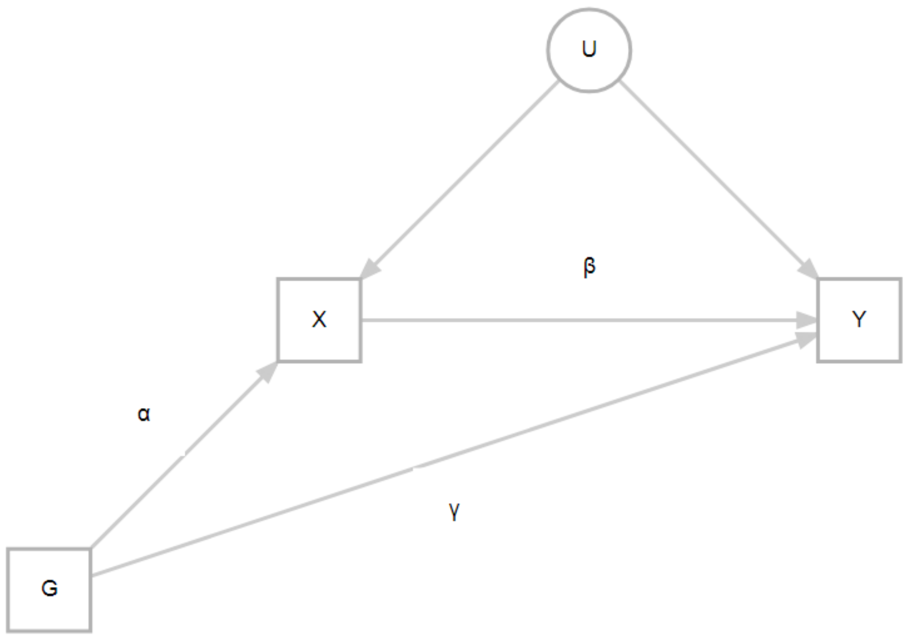

***Methods of MR Analysis***

We consider 5 two-sample methods of analysis:

- Fixed-effect IVW (IVW FE): The G-X and G-Y estimates are first obtained for each SNP; SNP-specific MR estimates are derived using the Wald estimator (ratio of G-Y over G-X), with standard error obtained using the Delta method [1]; the causal effect is then obtained by pooling SNP-specific MR estimates using a fixed-effect or a random-effects model, respectively. IVW FE is the most powerful two-sample MR method, but it assumes no pleiotropy [2].

- Random-effects IVW (IVW RE): It is performed in the same way as IVW FE, but a *multiplicative* random-effects meta-analysis model is used instead of a fixed-effect model to allow for pleiotropy. This method assumes that pleiotropic effects across SNPs are random (balanced pleiotropy), and that their magnitude is independent of the magnitude of the corresponding G-X effects (InSIDE assumption) [2]. Compared with an additive RE model, the multiplicative RE model is more robust to weak instrument bias since it downweighs variants with a weaker G-X association, and to the presence of outlying estimates, which are more likely to represent pleiotropic variants [2]. For these reasons, a multiplicative RE IVW has been recommended [3].

- Weighted median estimator: It assumes that more than 50% of the information contributing to the analysis comes from genetic variants that are valid (i.e. not pleiotropic) [4].

- Weighted mode estimator: It assumes that the largest weighted contribution of similar (identical in infinite samples) SNP-specific MR estimates comes from valid instruments [5].

- MR-Egger regression: G-Y estimates for the individual SNPs are regressed on their G-X estimates, with the intercept representing the overall pleiotropy and the slope the adjusted MR estimate [6]. MR-Egger makes the assumption of overall directional pleiotropy as well as the InSIDE assumption, and it works well only in the presence of a large spread of strengths, represented by the heterogeneity in G-X estimates across SNPs, $I^2_{GX}$, with a recommended $I^2_{GX}$ exceeding 90% [7].

In the one-sample MR, we also consider the 2SLS using individual-level data since this is a gold standard in the absence of pleiotropy.

***Parameters monitored***

For each method of analysis, for all scenarios and for both one-sample and two-sample data, we monitored the MR estimate and its standard error, as well as coverage and root mean square error (RMSE).

We also monitored:

- The ordinary least squares (OLS) estimate
- $R^2$ for the regression of X on all SNPs
- $R^2$ for the regression of Y on all SNPs
- $I^2$ for the G-X estimates
- An estimate of the residual correlation under the assumption of no pleiotropy
- Average SNP-specific F statistic
- Number of SNP-specific F-statistics below 10
- $I^2$ and Q for the FE IVW meta-analysis of SNP-specific MR estimates

The **residual correlation**, that is the correlation due to confounding rather than to the causal effect of X on Y, was estimated using the following approach: X is regressed on G, and fitted values ($f_X$) and residuals ($r_X$) are calculated; Y is then regressed on $f_X$ and the residuals are calculated ($r_Y$); the residual correlation due to confounding is estimated as the correlation between $r_X$ and ($r_Y$-beta*$r_X$), where beta is the MR estimate. This approach provided estimates very close to the true values in our simulations (data not shown), and can be used in practice in a MR study to estimate the correlation due to confounding.

The **F statistic**, a function of magnitude and precision of the G-X estimate that represents the strength of the SNP as an instrument for X [8], was calculated using the formula:
$F = GX^2/SE_{GX}^2$.

***Simulation parameters***

The simulation is based on a sample size of 300,000 and 100 independent SNPs with random allele frequency uniformly distributed between 1% and 99%.

The five correlations between the error components in X and Y (-0.4, -0.2, 0, 0.2, 0.4) represent different degrees of negative and positive confounding.

The true causal effect is either 0 (null) or 1.

SNP coefficients for the strength of association with X are exponential, so that there are many SNPs with small effects and a few with large effects. The figure below shows a histogram of the SNP coefficients for 1,000 simulated SNPs.

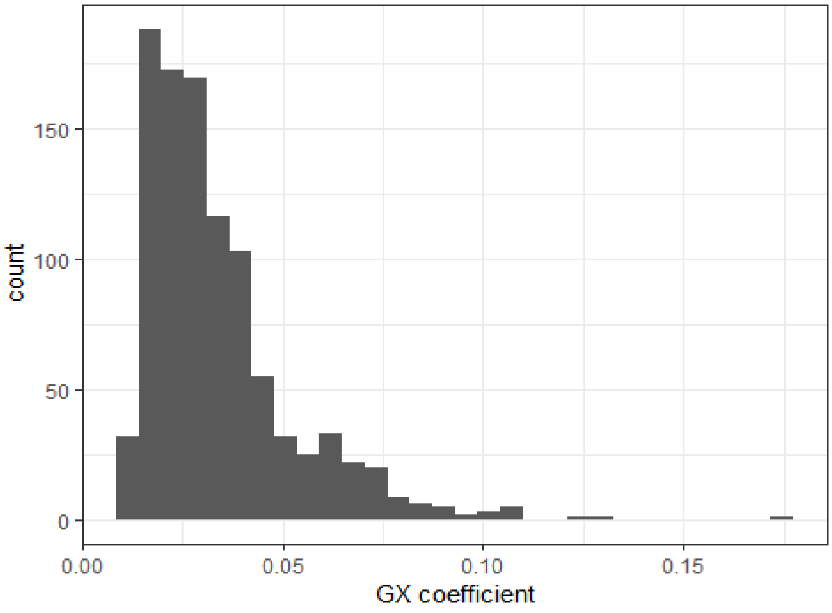


The SNP coefficients are adjusted so that rare SNPs cannot have very small coefficients, as otherwise the simulation would include large numbers of weak SNPs. The figure below shows the relationship between allele frequency and SNP coefficient for 1,000 simulated SNPs.

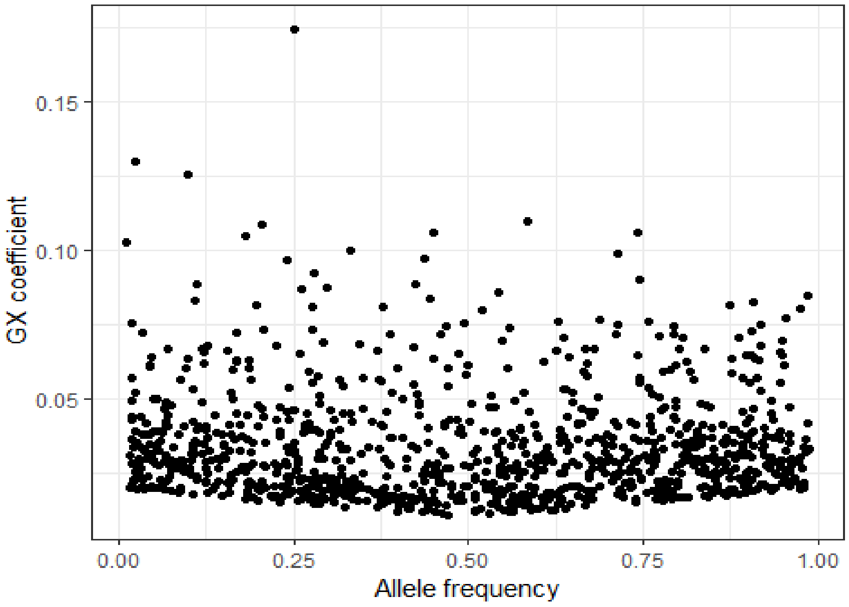


The individual SNPs have an average F-statistic of 67.9; out of each set of 100 SNPs, on average 5 SNPs had an F-statistic below 10 and would generally be considered as weak. SNP strength does not vary across the simulation experiment.

The average SNP explains 2.26% of the variance in X, and 0.22% of the variance in Y. The average percent of variance explained for G-X is the same for all scenarios, while for G-Y is larger when there is a causal relationship between X and Y and when there is a direct pleiotropic effect of G on Y.

Finally, the average $I^2$ is 90.8%, and this does not vary across the simulation experiment.

***Pleiotropy scenarios***

We generated data under three scenarios; no pleiotropy, directional pleiotropy in 20% of the SNPs, and balanced pleiotropy in 20% of the SNPs.

In the **directional pleiotropy** simulations, 20 of the 100 SNPs were pleiotropic in the sense of having a direct association between G and Y that did not pass through X. The coefficients for the pleiotropic effects were randomly chosen from the same distribution as was used for the G-X coefficients. Since both the SNP GX coefficients and the pleiotropic GY coefficients are positive, the pleiotropy will act to make the MR estimates larger. The bias of 0.15 to 0.19 across all scenarios with directional pleiotropy (with and without a causal effect, and with different degrees of correlation) observed for 2SLS, IVW FE and IVW RE (Supplementary table 4) gives an indication of the amount of pleiotropy that we have introduced by our choice of parameter values.

In the **balanced pleiotropy** simulations, 20 of the 100 SNPs were pleiotropic in the sense of having a direct association between G and Y that did not pass through X. The coefficients for the pleiotropic effects were randomly chosen from the same distribution as was used for the G-X coefficients, but then given a random sign so that on average the pleiotropy was zero.

**Supplementary Figure 1.** *Simulations with no causal effect in the presence of pleiotropy*: Q-Q plot of MR-Egger null p-values for different levels of correlation between X and Y due to confounding, in one-sample and two-sample MR. a) Balanced pleiotropy; b) Directional pleiotropy.

### a) Balanced pleiotropy

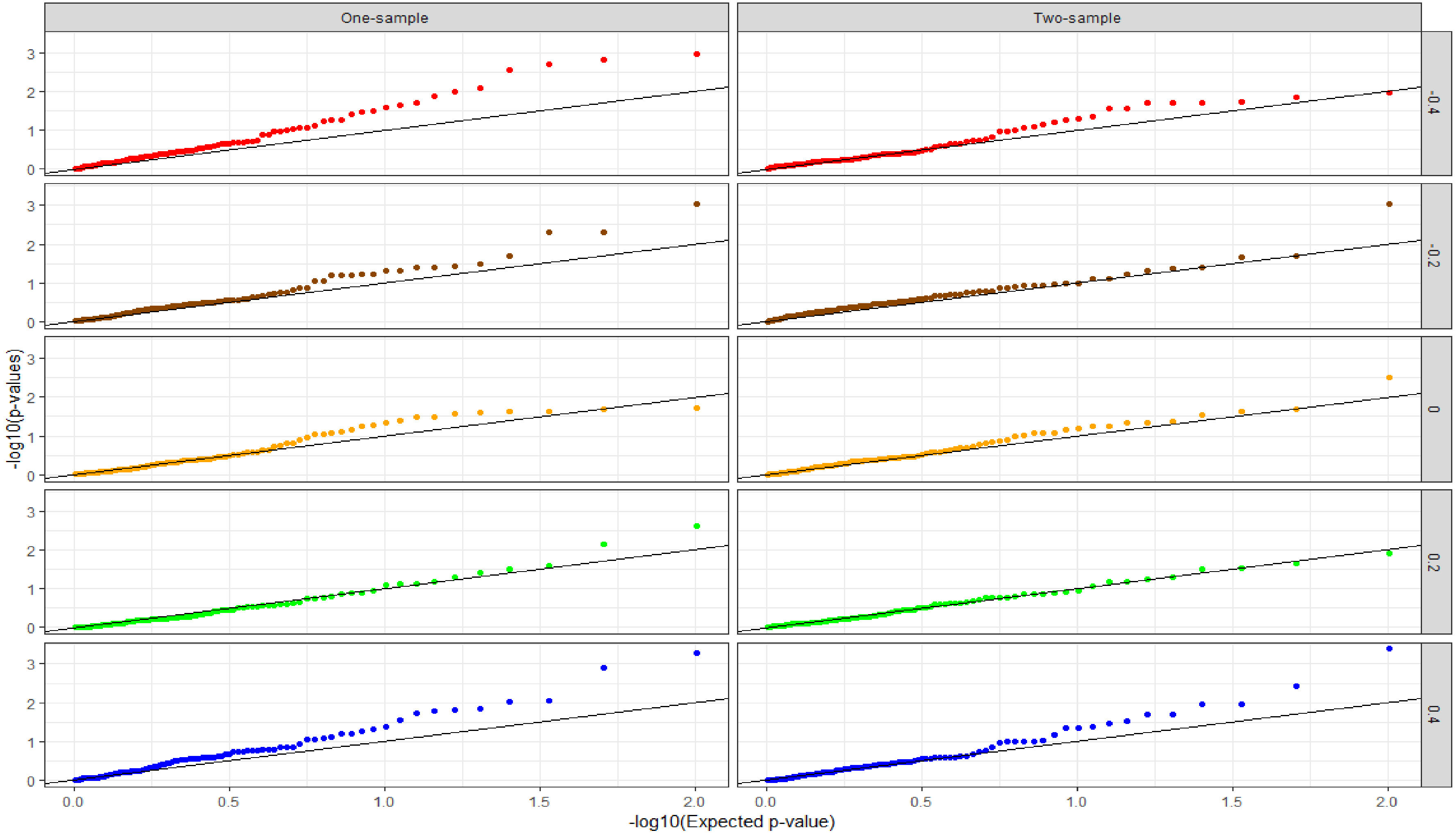

**b) Directional pleiotropy**

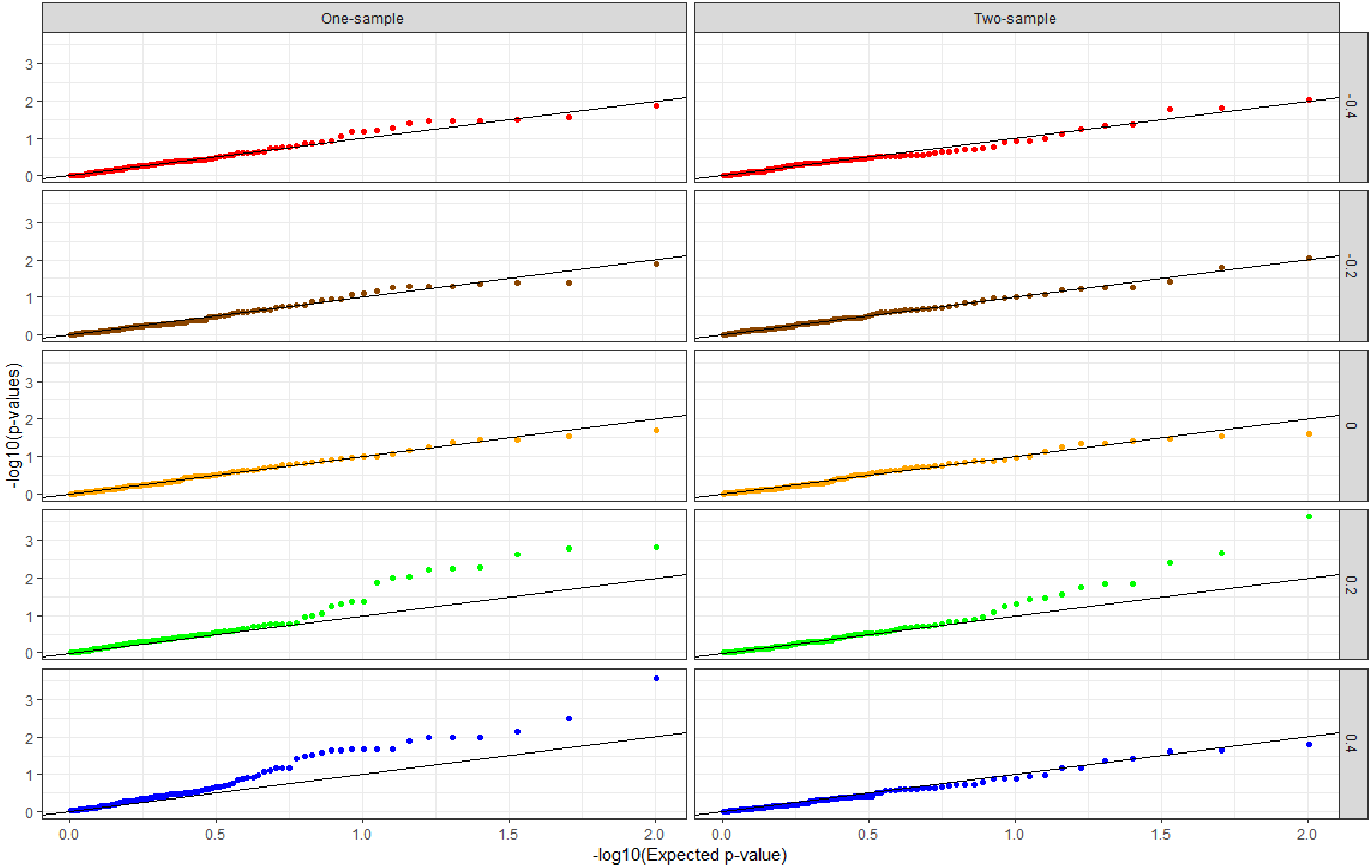

**Supplementary Figure 2.** Results of secondary analyses where the simulated G-X effect sizes have larger variability ($I^2_{GX}$ of 0.97). Outliers have been excluded to improve readability of the plots. a) No causal effect; b) Causal effect of 1.

**a) No causal effect**

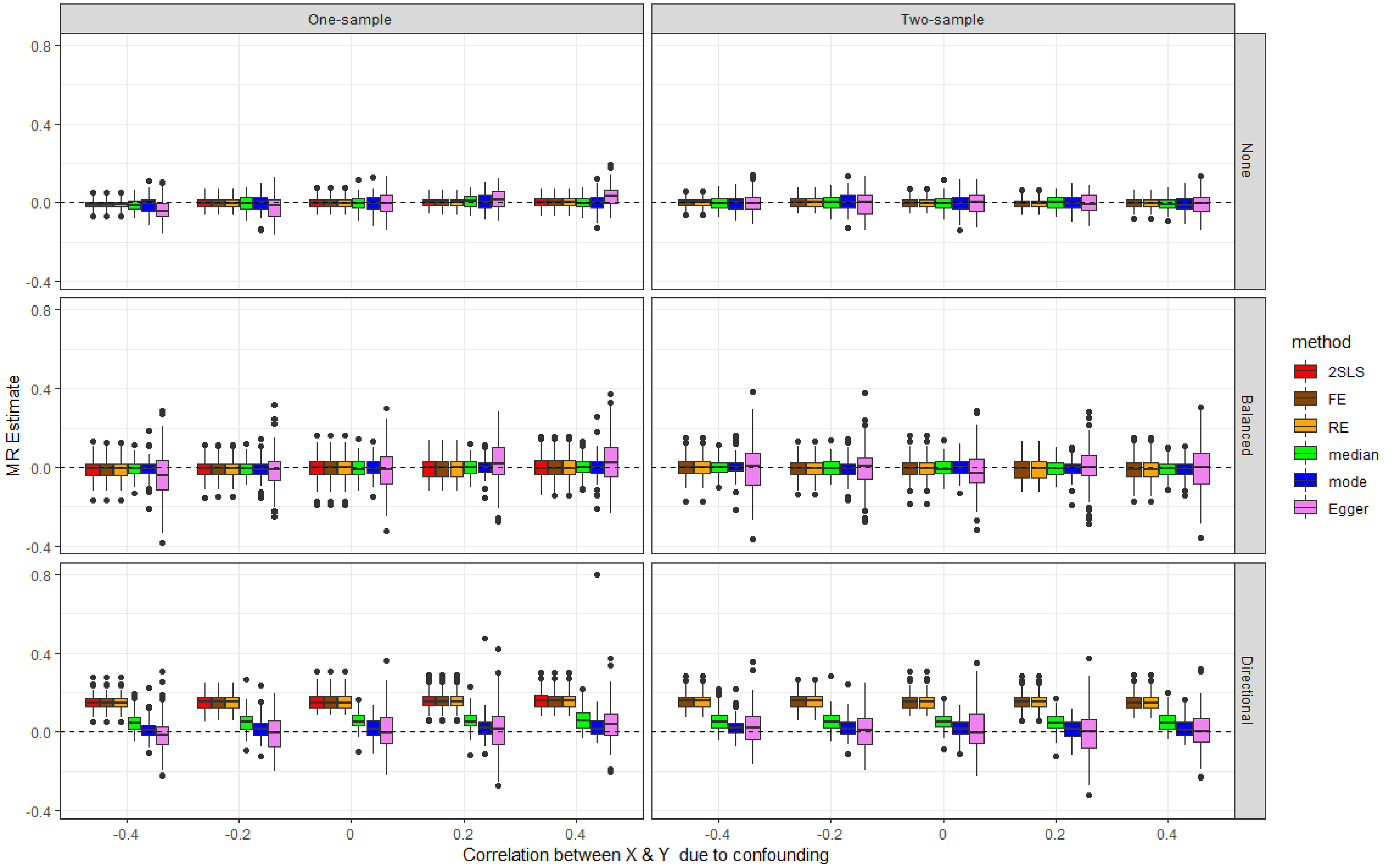

**b) Causal effect of 1**

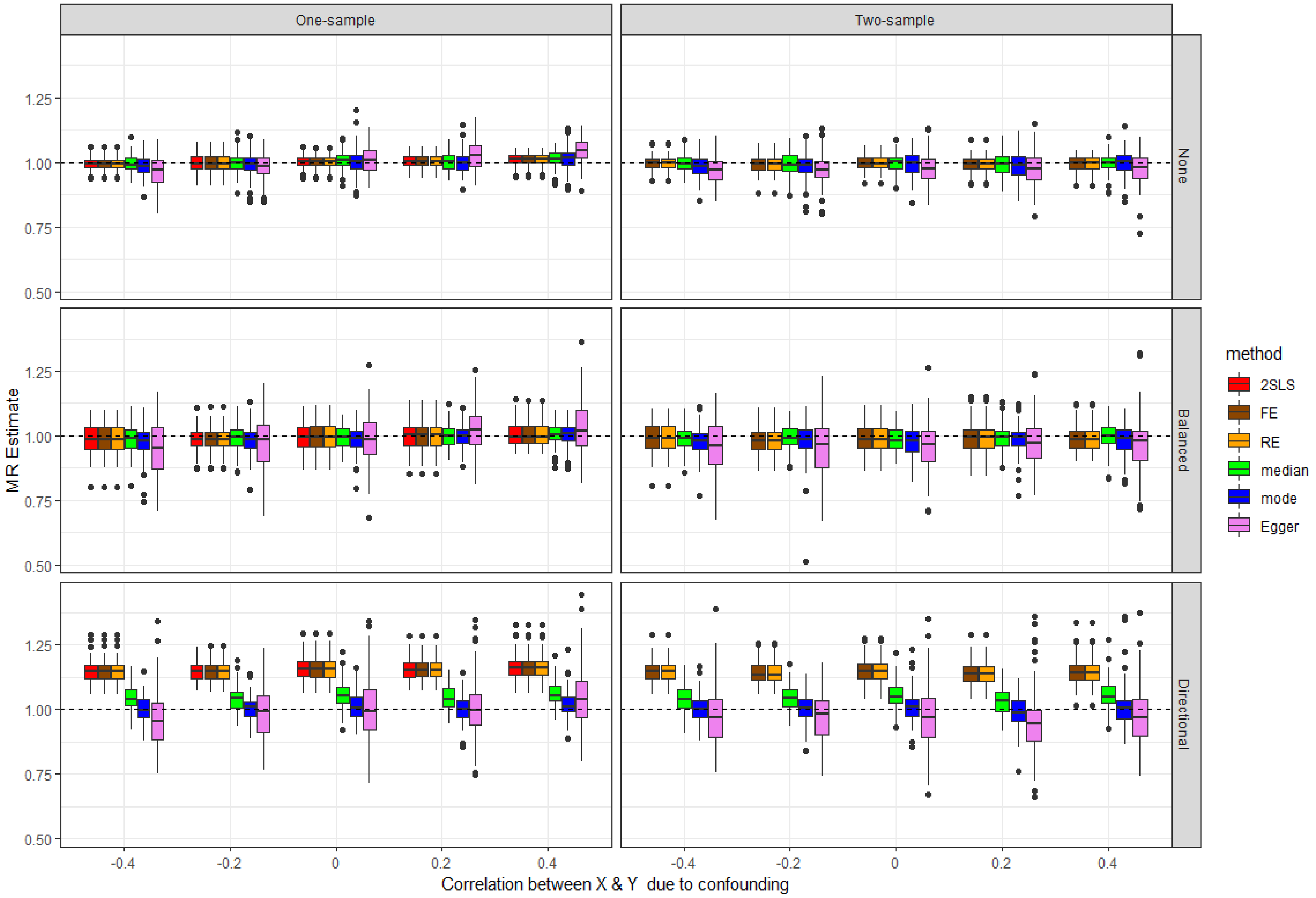

**Supplementary Table 1:** Characteristics and methods used in 27 MR investigations reported in the 10 papers reviewed. We only included one-sample MR analyses performed within UKB; when MR investigations were performed for multiple outcomes, we only considered one outcome per paper. IVW: inverse-variance weighted method; in brackets is specified whether a fixed-effect ("Fixed") or a random-effects ("Random") meta-analysis model was used, whenever this was specified. GRS: Genetic risk score. NR: Not reported. [a] Multiple exposures; [b] Overall F statistic for the GRS; [c] Binary exposures, for which % variance explained cannot be meaningfully estimated; [d] Multiple exposures and different population subgroups (males/females).

| Study | Sample size | N. SNPs | % Variance explained | F statistic | MR methods | |
|---|---|---|---|---|---|---|
| | | | | | Main analysis | Secondary analyses |
| Hindy, 2019 [9] | 376,435 | 76 | 7.3 | NR | IVW | MR-Egger, Weighted median estimator, Multivariable MR |
| Tyrrell, 2019 [10] | 287,503 | 14; 73 [a] | 0.2; 1.7 [a] | 86.5; 4,705 [b] | 2SLS using a GRS | IVW, MR-Egger, Median estimator, Penalized weighted median estimator |
| Taylor, 2019 [11] | 335,937 | 97 | NR | NR | IVW (Random) | MR-Egger, Weighted median estimator, Weighted mode estimator |
| Sun, 2019 [12] | 366,385 | 73 | 1.6 | 5,964 [b] | Ratio method using a GRS | MR-Egger, Weighted median estimator |
| Sun (2), 2019 [13] | 318,664 | 134; 233 [a] | / [c] | NR | IVW, MR-Egger, Simple median estimator, Weighted median estimator, MR-RAPS, MR-PRESSO | Leave-one-out |
| Khandaker, 2019 [14] | 367,703 | 2 to 185 [a] | NR | NR | IVW | MR-Egger, Weighted median estimator |
| Gharahkhani, 2019 [15] | 310,793 | 520 | 7 | NR | IVW (Fixed) | MR-Egger |
| Doherty, 2018 [16] | 278,374 | 24 to 68 [a] | NR | NR | Maximum likelihood MR | MR-Egger, Weighted median estimator, Weighted mode estimator, Leave-one-out, MR Steiger filtering |
| Wade, 2018 [17] | 335,308 | 77 | 1.82 | NR | Ratio method using a GRS | IVW, MR-Egger, Weighed median estimator, Weighted mode estimator |
| Minelli, 2018 [18] | 180,957 to 202,567 [d] | 24 to 206 [d] | / [c] | 10.1 to 382.5 [d] | IVW (Fixed) | IVW (Random), MR-Egger, Weighted median estimator, 2SLS |

**Supplementary Table 2.** *Simulations with **no pleiotropy***: Beta: causal effect estimate; SE: standard error; Cover.: coverage; RMSE: root mean square error.

| Data | Causal Effect | Confounding | 2SLS | | IVW FE | | IVW RE | | Median | | Mode | | Egger | |
|---|---|---|---|---|---|---|---|---|---|---|---|---|---|---|
| | | | **Beta<br>SE** | **Cover.<br>RMSE** | **Beta<br>SE** | **Cover.<br>RMSE** | **Beta<br>SE** | **Cover.<br>RMSE** | **Beta<br>SE** | **Cover.<br>RMSE** | **Beta<br>SE** | **Cover.<br>RMSE** | **Beta<br>SE** | **Cover.<br>RMSE** |
| **One-sample** | **0.0** | **-0.4** | -0.015<br>0.036 | 92.000<br>0.040 | -0.015<br>0.036 | 93.000<br>0.040 | -0.015<br>0.037 | 93.000<br>0.040 | -0.025<br>0.056 | 93.000<br>0.059 | -0.025<br>0.131 | 96.000<br>0.083 | -0.115<br>0.092 | 75.000<br>0.161 |
| | | **-0.2** | -0.014<br>0.037 | 93.000<br>0.037 | -0.015<br>0.037 | 93.000<br>0.037 | -0.015<br>0.038 | 94.000<br>0.037 | -0.017<br>0.057 | 97.000<br>0.051 | -0.019<br>0.088 | 97.000<br>0.072 | -0.055<br>0.096 | 86.000<br>0.121 |
| | | **0.0** | -0.000<br>0.036 | 95.000<br>0.034 | -0.001<br>0.036 | 95.000<br>0.034 | -0.001<br>0.037 | 95.000<br>0.034 | -0.002<br>0.056 | 98.000<br>0.048 | -0.002<br>0.089 | 100.000<br>0.064 | 0.002<br>0.093 | 96.000<br>0.087 |
| | | **0.2** | 0.015<br>0.036 | 88.000<br>0.044 | 0.015<br>0.036 | 88.000<br>0.044 | 0.015<br>0.037 | 89.000<br>0.044 | 0.019<br>0.056 | 95.000<br>0.052 | 0.019<br>0.087 | 98.000<br>0.069 | 0.062<br>0.092 | 87.000<br>0.115 |
| | | **0.4** | 0.017<br>0.036 | 89.000<br>0.039 | 0.017<br>0.036 | 89.000<br>0.039 | 0.017<br>0.037 | 90.000<br>0.039 | 0.022<br>0.057 | 96.000<br>0.051 | 0.026<br>0.083 | 98.000<br>0.066 | 0.108<br>0.092 | 83.000<br>0.137 |
| | **1.0** | **-0.4** | 0.986<br>0.036 | 95.000<br>0.038 | 0.986<br>0.033 | 92.000<br>0.038 | 0.986<br>0.036 | 96.000<br>0.038 | 0.979<br>0.056 | 95.000<br>0.052 | 0.965<br>0.124 | 95.000<br>0.072 | 0.889<br>0.088 | 80.000<br>0.147 |
| | | **-0.2** | 0.994<br>0.036 | 90.000<br>0.040 | 0.994<br>0.036 | 90.000<br>0.040 | 0.994<br>0.037 | 91.000<br>0.040 | 0.988<br>0.058 | 96.000<br>0.054 | 0.974<br>0.087 | 98.000<br>0.077 | 0.940<br>0.091 | 88.000<br>0.112 |
| | | **0.0** | 0.999<br>0.036 | 92.000<br>0.036 | 0.999<br>0.038 | 95.000<br>0.036 | 0.999<br>0.039 | 95.000<br>0.036 | 1.000<br>0.061 | 100.000<br>0.043 | 0.990<br>0.132 | 99.000<br>0.063 | 1.007<br>0.096 | 98.000<br>0.084 |
| | | **0.2** | 1.008<br>0.037 | 96.000<br>0.036 | 1.008<br>0.041 | 98.000<br>0.036 | 1.008<br>0.041 | 98.000<br>0.036 | 1.011<br>0.065 | 98.000<br>0.054 | 1.000<br>0.120 | 99.000<br>0.081 | 1.052<br>0.102 | 92.000<br>0.105 |
| | | **0.4** | 1.013<br>0.036 | 94.000<br>0.038 | 1.013<br>0.042 | 97.000<br>0.038 | 1.013<br>0.042 | 97.000<br>0.038 | 1.017<br>0.066 | 100.000<br>0.054 | 1.010<br>0.096 | 99.000<br>0.075 | 1.107<br>0.104 | 84.000<br>0.139 |

| Data | Causal Effect | Confounding | 2SLS | | IVW FE | | IVW RE | | Median | | Mode | | Egger | |
|---|---|---|---|---|---|---|---|---|---|---|---|---|---|---|
| | | | **Beta<br>SE** | **Cover.<br>RMSE** | **Beta<br>SE** | **Cover.<br>RMSE** | **Beta<br>SE** | **Cover.<br>RMSE** | **Beta<br>SE** | **Cover.<br>RMSE** | **Beta<br>SE** | **Cover.<br>RMSE** | **Beta<br>SE** | **Cover.<br>RMSE** |
| **Two-sample** | **0.0** | **-0.4** | | | 0.002<br>0.036 | 93.000<br>0.036 | 0.002<br>0.037 | 93.000<br>0.036 | -0.005<br>0.056 | 97.000<br>0.050 | -0.010<br>0.085 | 98.000<br>0.073 | -0.009<br>0.093 | 96.000<br>0.094 |
| | | **-0.2** | | | -0.005<br>0.037 | 96.000<br>0.036 | -0.005<br>0.038 | 96.000<br>0.036 | -0.004<br>0.057 | 100.000<br>0.052 | -0.005<br>0.157 | 99.000<br>0.070 | 0.003<br>0.097 | 90.000<br>0.108 |
| | | **0.0** | | | 0.000<br>0.036 | 94.000<br>0.034 | 0.000<br>0.037 | 94.000<br>0.034 | 0.001<br>0.056 | 98.000<br>0.048 | 0.002<br>0.088 | 99.000<br>0.066 | 0.008<br>0.093 | 96.000<br>0.090 |
| | | **0.2** | | | 0.006<br>0.036 | 90.000<br>0.040 | 0.006<br>0.037 | 90.000<br>0.040 | 0.005<br>0.056 | 96.000<br>0.048 | 0.004<br>0.083 | 97.000<br>0.069 | 0.006<br>0.091 | 96.000<br>0.090 |
| | | **0.4** | | | -0.000<br>0.036 | 95.000<br>0.035 | -0.000<br>0.037 | 96.000<br>0.035 | 0.000<br>0.057 | 100.000<br>0.044 | 0.003<br>0.124 | 100.000<br>0.083 | 0.005<br>0.092 | 96.000<br>0.084 |
| | **1.0** | **-0.4** | | | 0.986<br>0.033 | 94.000<br>0.038 | 0.986<br>0.036 | 98.000<br>0.038 | 0.978<br>0.055 | 97.000<br>0.051 | 0.966<br>0.081 | 98.000<br>0.071 | 0.904<br>0.087 | 82.000<br>0.133 |
| | | **-0.2** | | | 0.989<br>0.036 | 90.000<br>0.043 | 0.989<br>0.038 | 93.000<br>0.043 | 0.978<br>0.059 | 99.000<br>0.056 | 0.957<br>0.128 | 95.000<br>0.086 | 0.903<br>0.093 | 80.000<br>0.137 |
| | | **0.0** | | | 0.983<br>0.038 | 92.000<br>0.043 | 0.983<br>0.041 | 94.000<br>0.043 | 0.981<br>0.062 | 98.000<br>0.051 | 0.966<br>0.099 | 99.000<br>0.072 | 0.906<br>0.101 | 89.000<br>0.131 |
| | | **0.2** | | | 0.985<br>0.041 | 96.000<br>0.044 | 0.985<br>0.043 | 97.000<br>0.044 | 0.984<br>0.066 | 98.000<br>0.060 | 0.974<br>0.102 | 95.000<br>0.084 | 0.902<br>0.107 | 92.000<br>0.144 |
| | | **0.4** | | | 0.981<br>0.042 | 90.000<br>0.049 | 0.981<br>0.045 | 93.000<br>0.049 | 0.980<br>0.069 | 98.000<br>0.055 | 0.968<br>0.101 | 98.000<br>0.084 | 0.904<br>0.111 | 86.000<br>0.144 |

**Supplementary Table 3.** *Simulations with* ***balanced pleiotropy***: Beta: causal effect estimate; SE: standard error; Cover.: coverage; RMSE: root mean square error.

| Data | Causal Effect | Confounding | 2SLS | | IVW FE | | IVW RE | | Median | | Mode | | Egger | |
|---|---|---|---|---|---|---|---|---|---|---|---|---|---|---|
| | | | Beta<br>SE | Cover.<br>RMSE | Beta<br>SE | Cover.<br>RMSE | Beta<br>SE | Cover.<br>RMSE | Beta<br>SE | Cover.<br>RMSE | Beta<br>SE | Cover.<br>RMSE | Beta<br>SE | Cover.<br>RMSE |
| **One-sample** | **0.0** | **-0.4** | -0.017 | 75.500 | -0.017 | 75.600 | -0.017 | 93.000 | -0.022 | 94.000 | -0.025 | 96.400 | -0.107 | 86.700 |
| | | | 0.036 | 0.061 | 0.036 | 0.061 | 0.057 | 0.061 | 0.060 | 0.063 | 0.104 | 0.080 | 0.141 | 0.182 |
| | | **-0.2** | -0.010 | 77.100 | -0.010 | 76.800 | -0.010 | 94.900 | -0.012 | 93.900 | -0.012 | 96.200 | -0.055 | 91.400 |
| | | | 0.036 | 0.058 | 0.036 | 0.058 | 0.057 | 0.058 | 0.060 | 0.063 | 0.126 | 0.085 | 0.141 | 0.162 |
| | | **0.0** | -0.002 | 76.000 | -0.002 | 76.400 | -0.002 | 92.500 | -0.001 | 93.800 | -0.003 | 96.700 | -0.001 | 94.600 |
| | | | 0.036 | 0.061 | 0.036 | 0.061 | 0.057 | 0.061 | 0.060 | 0.064 | 0.114 | 0.085 | 0.142 | 0.145 |
| | | **0.2** | 0.008 | 74.700 | 0.008 | 74.900 | 0.008 | 94.000 | 0.011 | 93.700 | 0.013 | 96.700 | 0.057 | 91.800 |
| | | | 0.036 | 0.061 | 0.036 | 0.061 | 0.057 | 0.061 | 0.060 | 0.063 | 0.115 | 0.081 | 0.140 | 0.156 |
| | | **0.4** | 0.018 | 74.000 | 0.018 | 74.200 | 0.018 | 93.400 | 0.021 | 92.000 | 0.022 | 95.000 | 0.100 | 88.700 |
| | | | 0.036 | 0.061 | 0.036 | 0.061 | 0.057 | 0.061 | 0.060 | 0.065 | 0.109 | 0.083 | 0.141 | 0.177 |
| | **1.0** | **-0.4** | 0.983 | 73.200 | 0.983 | 69.500 | 0.983 | 93.400 | 0.976 | 92.400 | 0.956 | 92.000 | 0.890 | 87.400 |
| | | | 0.036 | 0.062 | 0.033 | 0.062 | 0.057 | 0.062 | 0.060 | 0.067 | 0.104 | 0.094 | 0.142 | 0.185 |
| | | **-0.2** | 0.991 | 76.400 | 0.991 | 75.600 | 0.991 | 92.600 | 0.989 | 94.100 | 0.973 | 95.500 | 0.952 | 92.100 |
| | | | 0.036 | 0.061 | 0.036 | 0.061 | 0.057 | 0.061 | 0.063 | 0.063 | 0.120 | 0.121 | 0.141 | 0.157 |
| | | **0.0** | 0.999 | 76.700 | 0.999 | 79.300 | 0.999 | 94.400 | 1.000 | 96.800 | 0.985 | 97.700 | 1.003 | 94.600 |
| | | | 0.036 | 0.059 | 0.038 | 0.059 | 0.057 | 0.059 | 0.065 | 0.060 | 0.125 | 0.081 | 0.142 | 0.147 |
| | | **0.2** | 1.008 | 75.700 | 1.008 | 80.300 | 1.008 | 92.000 | 1.013 | 97.600 | 1.001 | 98.200 | 1.057 | 95.000 |
| | | | 0.036 | 0.062 | 0.040 | 0.062 | 0.057 | 0.062 | 0.068 | 0.063 | 0.111 | 0.083 | 0.142 | 0.150 |
| | | **0.4** | 1.017 | 76.200 | 1.017 | 83.700 | 1.017 | 92.900 | 1.023 | 97.900 | 1.013 | 99.100 | 1.108 | 86.900 |
| | | | 0.036 | 0.061 | 0.042 | 0.061 | 0.057 | 0.061 | 0.071 | 0.062 | 0.142 | 0.087 | 0.141 | 0.183 |

| Data | Causal Effect | Confounding | 2SLS | | IVW FE | | IVW RE | | Median | | Mode | | Egger | |
|---|---|---|---|---|---|---|---|---|---|---|---|---|---|---|
| | | | Beta<br>SE | Cover.<br>RMSE | Beta<br>SE | Cover.<br>RMSE | Beta<br>SE | Cover.<br>RMSE | Beta<br>SE | Cover.<br>RMSE | Beta<br>SE | Cover.<br>RMSE | Beta<br>SE | Cover.<br>RMSE |
| **Two-sample** | **0.0** | **-0.4** | | | -0.000 | 75.900 | -0.000 | 95.100 | 0.001 | 95.400 | -0.000 | 97.000 | -0.006 | 94.100 |
| | | | | | 0.036 | 0.059 | 0.057 | 0.059 | 0.060 | 0.060 | 0.105 | 0.077 | 0.142 | 0.146 |
| | | **-0.2** | | | -0.002 | 77.700 | -0.002 | 94.900 | 0.001 | 94.300 | 0.004 | 96.600 | -0.002 | 93.200 |
| | | | | | 0.036 | 0.057 | 0.057 | 0.057 | 0.060 | 0.062 | 0.105 | 0.119 | 0.141 | 0.153 |
| | | **0.0** | | | -0.001 | 75.400 | -0.001 | 92.200 | -0.000 | 93.700 | -0.001 | 96.500 | 0.003 | 93.000 |
| | | | | | 0.036 | 0.062 | 0.057 | 0.062 | 0.060 | 0.064 | 0.111 | 0.085 | 0.142 | 0.151 |
| | | **0.2** | | | -0.001 | 76.300 | -0.001 | 94.100 | -0.002 | 94.600 | -0.001 | 97.300 | 0.002 | 94.600 |
| | | | | | 0.036 | 0.060 | 0.057 | 0.060 | 0.060 | 0.061 | 0.104 | 0.076 | 0.140 | 0.141 |
| | | **0.4** | | | 0.001 | 77.500 | 0.001 | 94.100 | -0.001 | 94.600 | -0.004 | 96.500 | -0.005 | 93.700 |
| | | | | | 0.036 | 0.059 | 0.057 | 0.059 | 0.060 | 0.061 | 0.114 | 0.081 | 0.141 | 0.145 |
| | **1.0** | **-0.4** | | | 0.986 | 70.200 | 0.986 | 92.600 | 0.980 | 92.900 | 0.959 | 93.900 | 0.906 | 88.400 |
| | | | | | 0.033 | 0.061 | 0.057 | 0.061 | 0.059 | 0.064 | 0.091 | 0.088 | 0.141 | 0.175 |
| | | **-0.2** | | | 0.985 | 73.300 | 0.985 | 92.600 | 0.979 | 93.800 | 0.960 | 94.700 | 0.915 | 89.300 |
| | | | | | 0.036 | 0.062 | 0.058 | 0.062 | 0.063 | 0.066 | 0.104 | 0.091 | 0.144 | 0.175 |
| | | **0.0** | | | 0.985 | 75.800 | 0.985 | 93.600 | 0.981 | 93.900 | 0.961 | 95.400 | 0.917 | 90.800 |
| | | | | | 0.038 | 0.063 | 0.059 | 0.063 | 0.066 | 0.069 | 0.123 | 0.095 | 0.148 | 0.175 |
| | | **0.2** | | | 0.986 | 77.500 | 0.986 | 92.200 | 0.983 | 94.700 | 0.965 | 95.000 | 0.916 | 91.900 |
| | | | | | 0.040 | 0.067 | 0.061 | 0.067 | 0.070 | 0.072 | 0.130 | 0.105 | 0.151 | 0.166 |
| | | **0.4** | | | 0.985 | 80.300 | 0.985 | 95.000 | 0.983 | 94.600 | 0.962 | 94.000 | 0.911 | 91.600 |
| | | | | | 0.042 | 0.065 | 0.062 | 0.065 | 0.073 | 0.073 | 0.144 | 0.109 | 0.154 | 0.181 |

**Supplementary Table 4.** *Simulations with* ***directional pleiotropy***: Beta: causal effect estimate; SE: standard error; Cover.: coverage; RMSE: root mean square error.

| Data | Causal Effect | Confounding | 2SLS | | IVW FE | | IVW RE | | Median | | Mode | | Egger | |
|---|---|---|---|---|---|---|---|---|---|---|---|---|---|---|
| | | | Beta<br>SE | Cover.<br>RMSE | Beta<br>SE | Cover.<br>RMSE | Beta<br>SE | Cover.<br>RMSE | Beta<br>SE | Cover.<br>RMSE | Beta<br>SE | Cover.<br>RMSE | Beta<br>SE | Cover.<br>RMSE |
| **One-sample** | **0.0** | **-0.4** | 0.150<br>0.037 | 3.700<br>0.157 | 0.150<br>0.036 | 3.600<br>0.157 | 0.150<br>0.055 | 15.700<br>0.157 | 0.056<br>0.061 | 85.600<br>0.082 | 0.005<br>0.106 | 97.800<br>0.075 | -0.094<br>0.134 | 89.800<br>0.163 |
| | | **-0.2** | 0.161<br>0.037 | 2.500<br>0.168 | 0.161<br>0.036 | 2.500<br>0.168 | 0.161<br>0.055 | 10.700<br>0.168 | 0.073<br>0.061 | 77.300<br>0.096 | 0.021<br>0.109 | 96.100<br>0.083 | -0.030<br>0.134 | 92.900<br>0.143 |
| | | **0.0** | 0.169<br>0.036 | 0.600<br>0.174 | 0.169<br>0.036 | 0.600<br>0.174 | 0.169<br>0.055 | 5.200<br>0.174 | 0.078<br>0.060 | 76.300<br>0.098 | 0.029<br>0.096 | 95.400<br>0.084 | 0.012<br>0.135 | 94.400<br>0.141 |
| | | **0.2** | 0.180<br>0.036 | 0.900<br>0.185 | 0.180<br>0.036 | 0.900<br>0.185 | 0.180<br>0.055 | 4.800<br>0.185 | 0.090<br>0.061 | 71.000<br>0.109 | 0.044<br>0.104 | 94.700<br>0.101 | 0.066<br>0.137 | 91.800<br>0.158 |
| | | **0.4** | 0.185<br>0.035 | 0.100<br>0.191 | 0.185<br>0.036 | 0.100<br>0.191 | 0.185<br>0.054 | 2.500<br>0.191 | 0.104<br>0.060 | 61.300<br>0.121 | 0.057<br>0.119 | 92.100<br>0.098 | 0.115<br>0.134 | 84.100<br>0.182 |
| | **1.0** | **-0.4** | 1.153<br>0.037 | 3.700<br>0.161 | 1.153<br>0.033 | 2.900<br>0.161 | 1.153<br>0.055 | 16.700<br>0.161 | 1.058<br>0.060 | 84.000<br>0.085 | 0.987<br>0.107 | 96.100<br>0.078 | 0.903<br>0.135 | 88.400<br>0.170 |
| | | **-0.2** | 1.160<br>0.037 | 2.800<br>0.167 | 1.160<br>0.036 | 2.700<br>0.167 | 1.160<br>0.055 | 11.600<br>0.167 | 1.068<br>0.063 | 84.100<br>0.092 | 1.004<br>0.106 | 97.200<br>0.091 | 0.960<br>0.134 | 93.500<br>0.145 |
| | | **0.0** | 1.167<br>0.036 | 0.700<br>0.173 | 1.167<br>0.038 | 1.300<br>0.173 | 1.167<br>0.055 | 7.800<br>0.173 | 1.079<br>0.066 | 80.300<br>0.100 | 1.012<br>0.111 | 98.400<br>0.131 | 1.005<br>0.136 | 94.100<br>0.138 |
| | | **0.2** | 1.175<br>0.036 | 0.400<br>0.181 | 1.175<br>0.040 | 1.200<br>0.181 | 1.175<br>0.054 | 5.500<br>0.181 | 1.090<br>0.069 | 77.500<br>0.109 | 1.028<br>0.114 | 97.300<br>0.085 | 1.062<br>0.134 | 90.400<br>0.154 |
| | | **0.4** | 1.187<br>0.036 | 0.400<br>0.192 | 1.187<br>0.043 | 0.900<br>0.192 | 1.187<br>0.054 | 2.600<br>0.192 | 1.105<br>0.072 | 72.200<br>0.120 | 1.042<br>0.116 | 98.000<br>0.086 | 1.123<br>0.136 | 84.200<br>0.186 |

| Data | Causal Effect | Confounding | 2SLS | | IVW FE | | IVW RE | | Median | | Mode | | Egger | |
|---|---|---|---|---|---|---|---|---|---|---|---|---|---|---|
| | | | Beta<br>SE | Cover.<br>RMSE | Beta<br>SE | Cover.<br>RMSE | Beta<br>SE | Cover.<br>RMSE | Beta<br>SE | Cover.<br>RMSE | Beta<br>SE | Cover.<br>RMSE | Beta<br>SE | Cover.<br>RMSE |
| **Two-sample** | **0.0** | **-0.4** | | | 0.167 | 0.900 | 0.167 | 7.900 | 0.078 | 76.600 | 0.030 | 96.300 | 0.009 | 95.800 |
| | | | | | 0.036 | 0.173 | 0.054 | 0.173 | 0.061 | 0.098 | 0.124 | 0.083 | 0.134 | 0.131 |
| | | **-0.2** | | | 0.169 | 1.200 | 0.169 | 7.600 | 0.084 | 71.000 | 0.035 | 94.600 | 0.021 | 94.200 |
| | | | | | 0.036 | 0.176 | 0.054 | 0.176 | 0.061 | 0.104 | 0.096 | 0.086 | 0.134 | 0.138 |
| | | **0.0** | | | 0.168 | 0.700 | 0.168 | 5.900 | 0.078 | 75.700 | 0.032 | 95.500 | 0.011 | 94.400 |
| | | | | | 0.036 | 0.174 | 0.055 | 0.174 | 0.060 | 0.098 | 0.120 | 0.146 | 0.135 | 0.141 |
| | | **0.2** | | | 0.171 | 1.400 | 0.171 | 7.400 | 0.079 | 76.300 | 0.032 | 96.200 | 0.013 | 93.200 |
| | | | | | 0.036 | 0.177 | 0.055 | 0.177 | 0.061 | 0.099 | 0.103 | 0.088 | 0.137 | 0.144 |
| | | **0.4** | | | 0.169 | 0.600 | 0.169 | 7.300 | 0.081 | 73.000 | 0.035 | 95.100 | 0.012 | 93.500 |
| | | | | | 0.036 | 0.175 | 0.055 | 0.175 | 0.061 | 0.103 | 0.105 | 0.087 | 0.135 | 0.141 |
| | **1.0** | **-0.4** | | | 1.155 | 2.800 | 1.155 | 14.700 | 1.058 | 84.700 | 0.991 | 96.300 | 0.918 | 88.400 |
| | | | | | 0.033 | 0.162 | 0.055 | 0.162 | 0.060 | 0.086 | 0.105 | 0.078 | 0.134 | 0.164 |
| | | **-0.2** | | | 1.154 | 4.100 | 1.154 | 16.600 | 1.063 | 84.600 | 0.997 | 95.900 | 0.922 | 91.400 |
| | | | | | 0.036 | 0.161 | 0.056 | 0.161 | 0.064 | 0.090 | 0.113 | 0.086 | 0.137 | 0.160 |
| | | **0.0** | | | 1.153 | 4.700 | 1.153 | 19.700 | 1.067 | 83.500 | 0.999 | 98.200 | 0.916 | 89.300 |
| | | | | | 0.038 | 0.161 | 0.058 | 0.161 | 0.067 | 0.094 | 0.135 | 0.085 | 0.142 | 0.172 |
| | | **0.2** | | | 1.151 | 7.000 | 1.151 | 24.600 | 1.067 | 84.900 | 1.003 | 96.700 | 0.918 | 90.600 |
| | | | | | 0.040 | 0.159 | 0.059 | 0.159 | 0.070 | 0.098 | 0.126 | 0.112 | 0.144 | 0.169 |
| | | **0.4** | | | 1.156 | 7.500 | 1.156 | 22.700 | 1.073 | 84.700 | 1.006 | 97.400 | 0.923 | 91.300 |
| | | | | | 0.043 | 0.164 | 0.061 | 0.164 | 0.074 | 0.102 | 0.136 | 0.094 | 0.150 | 0.173 |

**Supplementary Table 5.** *Ordinary least squares results for the regression analysis of Y on X in the one-sample data*. OLS: ordinary least squares; SE: standard error: RMSE: root mean square error.

| Pleiotropy | Causal Effect | Confounding | OLS Estimate | OLS SE | Bias | Coverage | RMSE |
|---|---|---|---|---|---|---|---|
| **None** | **0.0** | **-0.4** | -1.172 | 0.0050 | -1.172 | 0.0 | 1.172 |
| | | **-0.2** | -0.586 | 0.0053 | -0.586 | 0.0 | 0.586 |
| | | **0.0** | -0.000 | 0.0054 | -0.000 | 97.0 | 0.005 |
| | | **0.2** | 0.587 | 0.0053 | 0.587 | 0.0 | 0.587 |
| | | **0.4** | 1.173 | 0.0050 | 1.173 | 0.0 | 1.173 |
| | **1.0** | **-0.4** | -0.174 | 0.0050 | -1.174 | 0.0 | 1.174 |
| | | **-0.2** | 0.414 | 0.0053 | -0.586 | 0.0 | 0.586 |
| | | **0.0** | 0.999 | 0.0054 | -0.001 | 100.0 | 0.005 |
| | | **0.2** | 1.586 | 0.0053 | 0.586 | 0.0 | 0.586 |
| | | **0.4** | 2.174 | 0.0050 | 1.174 | 0.0 | 1.174 |
| **Balanced** | **0.0** | **-0.4** | -1.174 | 0.0050 | -1.174 | 0.0 | 1.174 |
| | | **-0.2** | -0.588 | 0.0053 | -0.588 | 0.0 | 0.588 |
| | | **0.0** | 0.001 | 0.0054 | 0.001 | 93.0 | 0.006 |
| | | **0.2** | 0.585 | 0.0053 | 0.585 | 0.0 | 0.585 |
| | | **0.4** | 1.173 | 0.0050 | 1.173 | 0.0 | 1.173 |
| | **1.0** | **-0.4** | -0.172 | 0.0050 | -1.172 | 0.0 | 1.172 |
| | | **-0.2** | 0.414 | 0.0053 | -0.586 | 0.0 | 0.586 |
| | | **0.0** | 1.001 | 0.0054 | 0.001 | 98.0 | 0.005 |
| | | **0.2** | 1.587 | 0.0053 | 0.587 | 0.0 | 0.587 |
| | | **0.4** | 2.174 | 0.0050 | 1.174 | 0.0 | 1.174 |
| **Directional** | **0.0** | **-0.4** | -1.170 | 0.0050 | -1.170 | 0.0 | 1.170 |
| | | **-0.2** | -0.582 | 0.0053 | -0.582 | 0.0 | 0.582 |
| | | **0.0** | 0.004 | 0.0054 | 0.004 | 86.0 | 0.007 |
| | | **0.2** | 0.591 | 0.0053 | 0.591 | 0.0 | 0.591 |
| | | **0.4** | 1.176 | 0.0050 | 1.176 | 0.0 | 1.176 |
| | **1.0** | **-0.4** | -0.169 | 0.0050 | -1.169 | 0.0 | 1.169 |
| | | **-0.2** | 0.417 | 0.0053 | -0.583 | 0.0 | 0.583 |
| | | **0.0** | 1.004 | 0.0054 | 0.004 | 85.0 | 0.007 |
| | | **0.2** | 1.590 | 0.0053 | 0.590 | 0.0 | 0.590 |
| | | **0.4** | 2.177 | 0.0050 | 1.177 | 0.0 | 1.177 |
